# An S1 subunit vaccine and combination adjuvant (COVAC-1) elicits robust protection against SARS-CoV-2 challenge in African green monkeys

**DOI:** 10.1101/2022.06.16.496375

**Authors:** Lauren Garnett, Kaylie N Tran, Mable Chan, Kevin Tierney, Zachary Schiffman, Jonathan Audet, Jocelyne M. Lew, Courtney Meilleur, Michael Chan, Kathy Manguiat, Nikesh Tailor, Robert Vendramelli, Yvon Deschambault, Guillaume Beaudoin-Bussières, Catherine Bourassa, Jonathan Richard, Andrés Finzi, Rick Higgins, Sylvia van Drunen Littlel-van den Hurk, Volker Gerdts, James E Strong, Darryl Falzarano

## Abstract

Severe acute respiratory syndrome coronavirus 2 (SARS-CoV-2) is the agent responsible for the ongoing global pandemic. With over 500 million cases and more than 6 million deaths reported globally, the need for access to effective vaccines is clear. An ideal SARS-CoV-2 vaccine will prevent pathology in the lungs and prevent virus replication in the upper respiratory tract, thus reducing transmission. Here, we assessed the efficacy of an adjuvanted SARS-CoV-2 S1 subunit vaccine, called COVAC-1, in an African green monkey (AGM) model. AGMs immunized and boosted with COVAC-1 were protected from SARS-CoV-2 challenge compared to unvaccinated controls based on reduced pathology and reduced viral RNA levels and infectious virus in the respiratory tract. Both neutralizing antibodies and antibodies capable of mediating antibody-dependent cell-mediated cytotoxicity (ADCC) were observed in vaccinated animals prior to the challenge. COVAC-1 induced effective protection, including in the upper respiratory tract, thus supporting further development and utility for determining the mechanism that confers this protection.

**AUTHOR SUMMARY:** Vaccines that can prevent the onward transmission of SARS-CoV-2 and prevent disease are highly desirable. Whether this can be accomplished without mucosal immunization by a parenterally administered subunit vaccine is not well established. Here we demonstrate that following two vaccinations, a protein subunit vaccine containing the S1 portion of the SARS-CoV-2 spike glycoprotein and the novel adjuvant TriAdj significantly reduces the amount of virus in the lungs and also mediates rapid clearance of the virus from the upper respiratory tract. Further support of the effectiveness of COVAC-1 was the observation of reduced pathology in the lungs and viral RNA being largely absent from tissues, blood, and rectal swabs. Thus COVAC-1 appears promising at mediating protection in both the upper and lower respiratory tract and may be capable of reducing subsequent transmission of SARS-CoV-2. Further investigation into the mechanism of protection in the upper respiratory tract and the initial immune response that supports this would be warranted.

## INTRODUCTION

Severe acute respiratory syndrome coronavirus 2 (SARS-CoV-2), the pathogen responsible for Coronavirus disease 2019 (COVID-19), has caused over 500 million cases and over 6 million deaths globally (WHO dashboard) as of May 2022. With the rising case numbers internationally and the inequality of vaccine accessibility throughout the world, the need to develop more effective vaccines is ongoing, particularly vaccines that can reduce transmission. Several vaccines have successfully passed phase 3 clinical trials and/or have gained regulatory approval and have been proven to elicit effective protection against SARS-CoV-2, with more candidates going through the developmental pipeline [1-4]. However, even with multiple vaccines approved, worldwide vaccine coverage remains too low. Furthermore, continued development of new vaccines is needed given the emergence of new variants of concern (VOC) that limit the efficacy of current vaccines, as well as hurdles demonstrated in development, clinical trials, production, and distribution.

The majority of vaccines against SARS-CoV-2, both approved and in development, include the spike (S) glycoprotein [5-8]. The trimeric S glycoprotein is made up of two subunits, S1 and S2. S is essential for mediating binding, fusion, and uptake of virions into mammalian cells, as well as being the target of neutralizing antibodies [9]. Of the various vaccine platforms, including nucleic acid, viral-vectored, inactivated, and protein subunit, recombinant protein subunit vaccines have some advantages over other vaccine platforms, including greater safety while reducing cost and handling restrictions [10]. Although subunit vaccines typically induce weaker neutralizing antibodies, studies on recombinant subunit vaccines that include the receptor binding domain (RBD) have shown higher neutralizing antibodies with no antibody-dependent enhancement (ADE) effects, suggesting both a safe and effective vaccine platform [11-13]. Furthermore, adjuvants may augment and prolong the immune output without corresponding increases in deleterious effects [14].

We aimed to determine whether an S1-containing subunit protein vaccine and a combination adjuvant platform developed at the Vaccine and Infectious Disease Organization (VIDO) elicits effective immunogenicity and protection against SARS-CoV-2 in African green monkeys (AGMs) [15-17]. The vaccine candidate, COVAC-1, contains a codon optimized, mammalian-produced S1 segment of the SARS-CoV-2 spike glycoprotein and an adjuvant, TriAdj. TriAdj is comprised of the toll-like receptor (TLR) agonist poly I-C, poly[di(sodium carboxylatoethylphenoxy)phosphazene] (PCEP), and the synthetic cationic peptide IDR-100. This combination adjuvant has been used in numerous animal species with no adverse reactions for vaccines with both viral and bacterial antigens, including one for human respiratory syncytial virus [18]. TriAdj has also been shown to induce a balanced or Th1-biased immune response that provides long-lasting immunity in various animal models, including mice, hamsters, cotton rats, sheep, alpacas and pigs [18-22]. This combination adjuvant has also frequently outperformed existing commercial adjuvants [23].

## RESULTS

### Clinical parameters are comparable between vaccinated and control animals

Six AGMs (Animal numbers 1478, 1512, 1540, 1572, 1775, and 1820) were vaccinated with COVAC-1, a mammalian-produced HIS-tagged SARS-CoV-2 S1 protein formulated with TriAdj adjuvant 56 days before challenge, followed by a homologous boost 28 days before challenge. Six control animals (1164, 1250, 1342, 1687, 1774, and 1776) received PBS and were used as comparators following the same timeline as the vaccine group. All twelve AGMs were challenged with a total of 1.54E+04 TCID_50_ of SARS-CoV-2 (Canada/ON-VIDO-01/2020) by a combination of intratracheal (i.t), intranasal (i.n), oral and intraocular (i.o) routes similar to that described by Blair et al. [24]. Longitudinal oral, rectal, and nasal sampling was performed along with blood and bronchoalveolar lavage (BAL) collections to assess viral load and shedding (Figure 1A). No animals had overt signs of clinical disease such as fever (Figure 1B), weight loss (Figure 1C), or changes in behaviour or appetite. However, there were trends towards increased pCO_2_ and bicarbonate following infection, especially in the control animals. This coincided with an increase in respiratory rate noted particularly in the control group at 3dpi with no substantial changes in oxygen saturation (Figure 2A-E).

**Figure 1:**
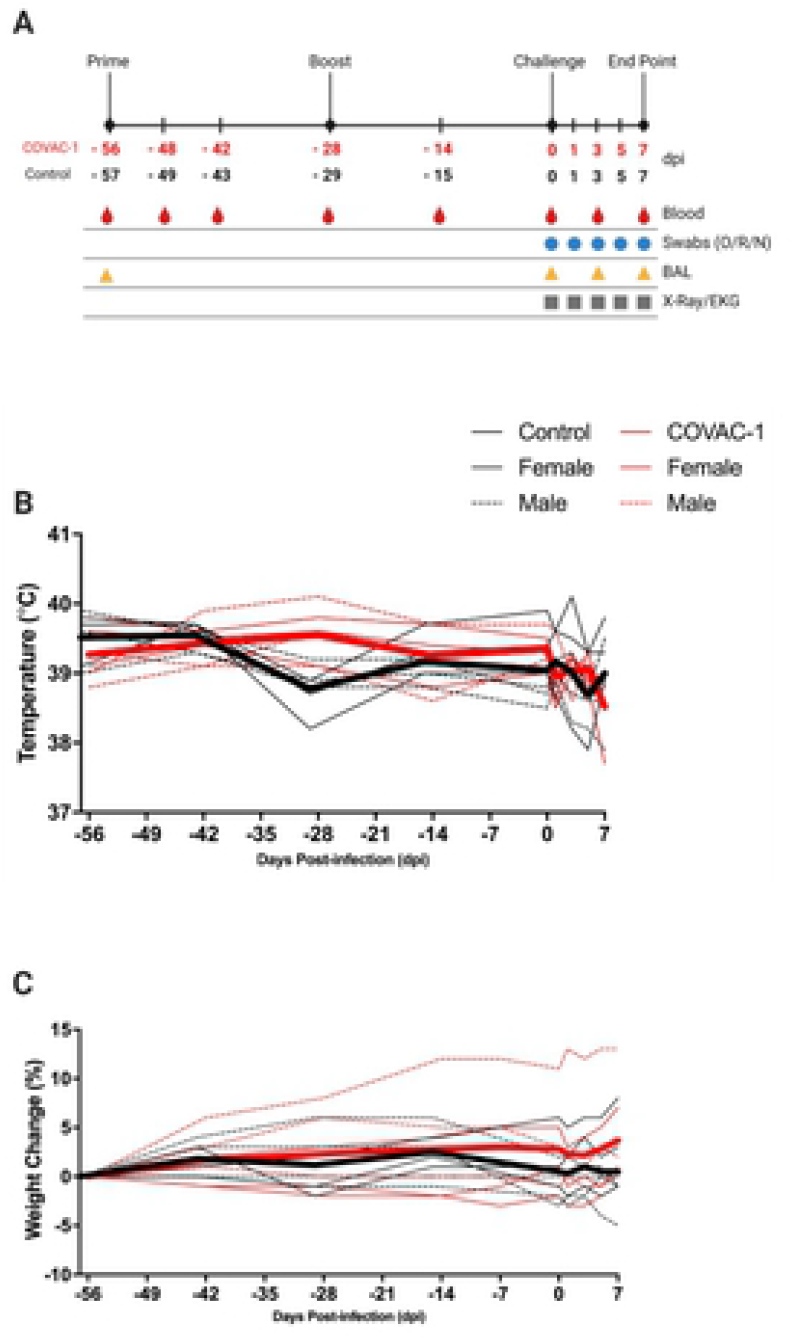
Schematic of the experimental and sampling timeline showing initial COVAC-1 vaccination and homologous boost prior to SARS-CoV-2 challenge, as well as the time points BAL, blood, X-ray/EKGs and swabs, taken throughout (A). All animals were monitored for changes in temperature (B) and percent weight change (C) from initial vaccination (day-56/-57) until 7dpi. Each line represents an individual animal, with red indicating vaccinated animals and black representing control AGMS.

**Figure 2:**
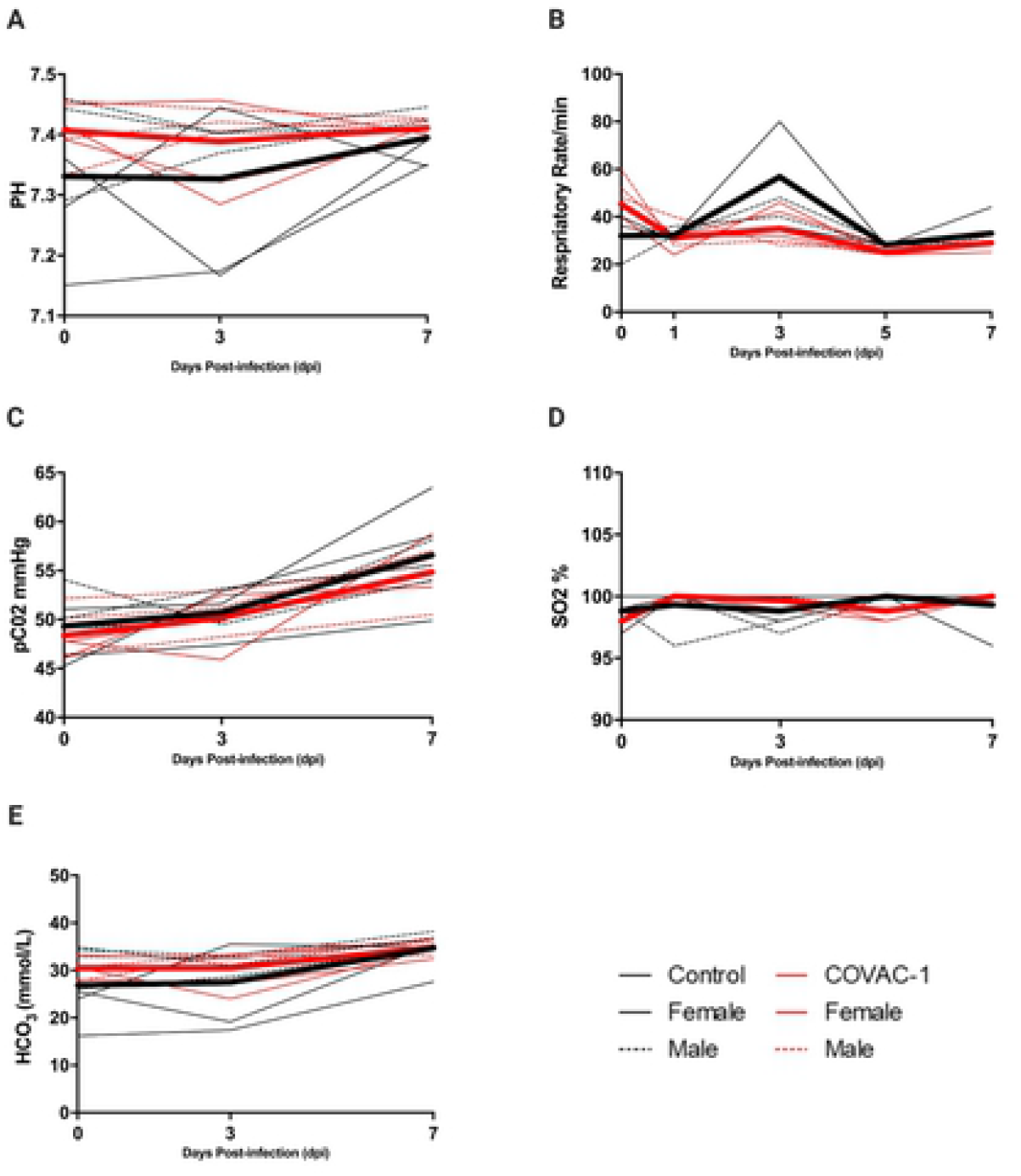
Blood was collected from all AGMs at 0, 3 and 7dpi for blood gas analysis using an iSTAT Alinity hematological analyzer. The measured parameters included pH (A), pCO_2_ (C), saturated oxygen sO_2_ (D) and bicarbonate HCO_3_ (E). Animals’ respiratory rates were measured during sedation (B). Each line represents an individual animal, with red indicating vaccinated animals and black representing control AGMs.

White blood cell (WBC) count, red blood cell (RBC) count, lymphocytes, platelets, monocytes, and neutrophils were all within the normal range prior to infection (Figure 3A-F). During the peak of infection (3dpi), counts of WBC, platelets and neutrophils decreased in both the COVAC-1 vaccinated animals and the control animals. These counts started to recover again by the end of the study (7dpi) (Figure 3A, B, F). Average alkaline phosphatase (ALP) and albumin (ALB), typically cited as markers for liver function, decreased slightly during infection. At the same time, aminotransferase (ALT), an enzyme signifying hepatocellular injury, was elevated in both the vaccinated and control groups (Figure 4A, C, E), but was decreasing towards normal in COVAC-1 vaccinated animals by 7dpi. Blood urea nitrogen (BUN) and creatinine (CRE) levels, measures of kidney function, were found to be within the normal range (Figure 4B, D). Total protein (TP) levels, a combined measurement of liver and kidney function, decreased from baseline at 7dpi for both vaccinated and control AGMs (Figure 4F). Thoracic radiographs and EKGs taken on days 0, 1, 3, 5, and 7 post-infection showed no changes over baseline measurements (data not shown).

**Figure 3:**
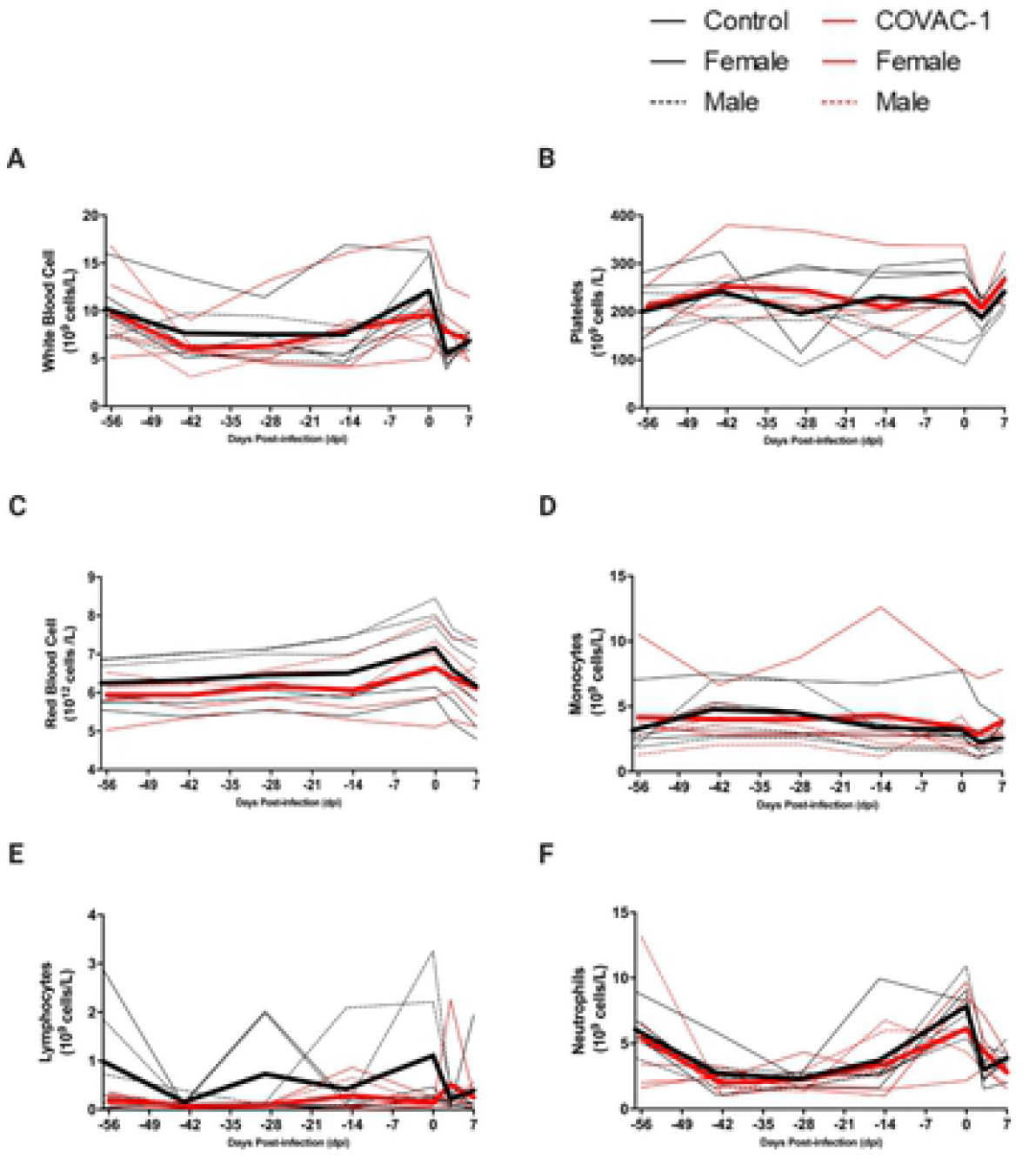
EDTA treated blood was collected from all AGMs at 0, 3, and 7 days post SARS-CoV-2 infection for complete blood count analysis. Hematology values are presented for total white blood cell counts (A), platelets (B), total red blood cell counts (C), monocyte (D), lymphocytes (E) and neutrophils (F). Each line represents an individual animal, with red indicating vaccinated animals and black representing control AGMS.

**Figure 4:**
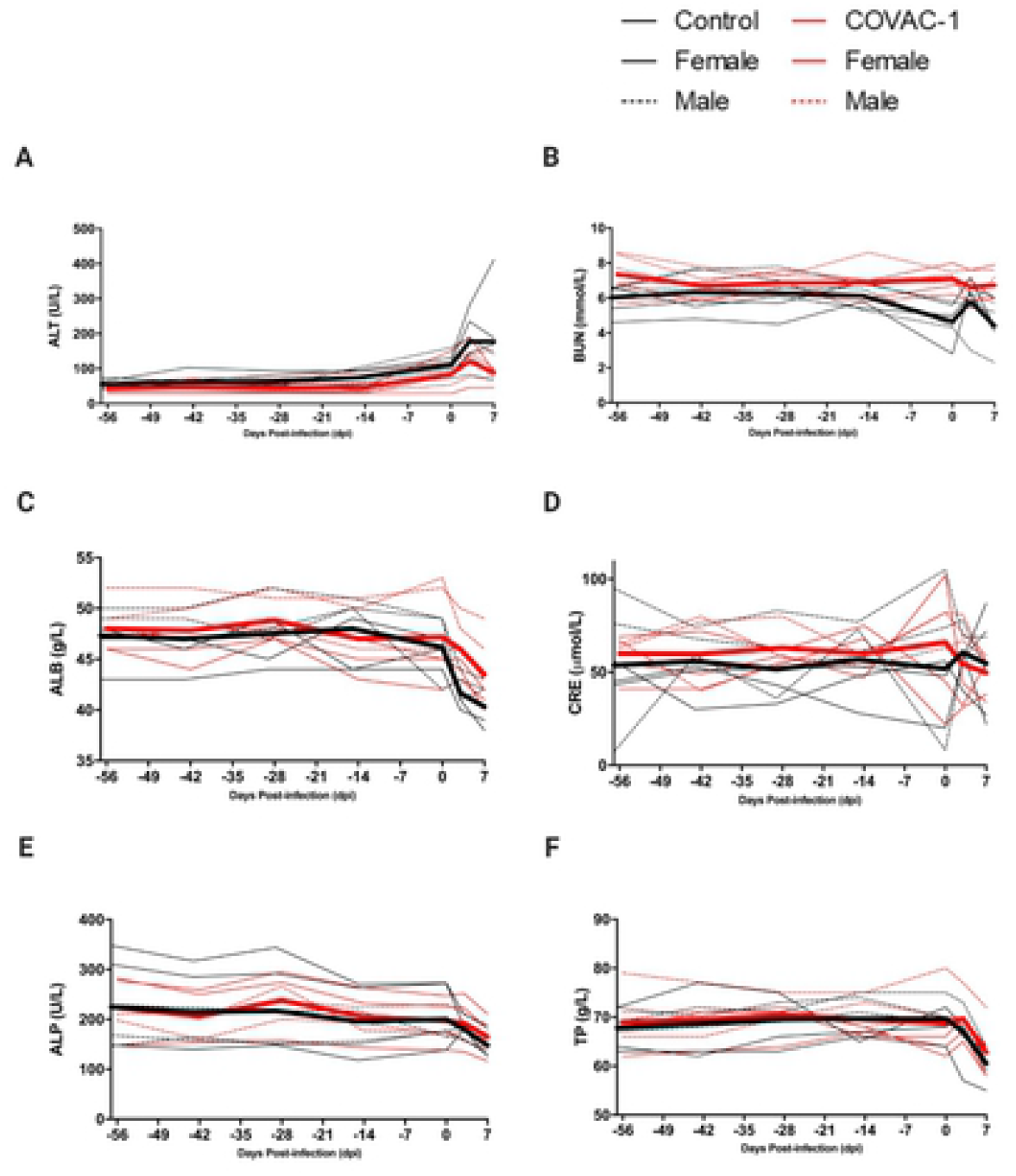
Lithium heparin treated blood collected at 0, 3 and 7 days post infection with SARS-CoV-2 was used to evaluate clinical chemistry markers for kidney and liver function. These markers include alanine aminotransferase ALT (A), blood urea nitrogen BUN (B), albumin ALB (C), creatinine CRE (D), alkaline phosphatase ALP (E) and total protein TP (F). Each line represents an individual animal, with red indicating vaccinated animals and black representing control AGMS.

### COVAC-1 reduces the quantity and duration of viral RNA and infectious virus

Viral loads and infectious virus were quantified from oral, rectal, and nasal swabs collected on days 0, 1, 3, 5 and 7 post-infection by RT-qPCR and TCID_50_ assay (Figure 5). On day 1 post-challenge, both vaccinated and unvaccinated animals had equivalent levels of viral RNA, of approximately 10^4^ genome equivalence (GEQ)/mL, in nasal swabs. Given the relatively high challenge dose, this was not unexpected. On subsequent days the level of viral RNA remained relatively constant in control animals. In contrast, vaccinated animals began to show substantially lower levels of viral RNA, with a 2.57 log reduction on day 3 (p=0.0025), a 2.80 log reduction on day 5 (p=0.016), and no detectable viral RNA by day 7 (Figure 5A). Infectious virus recovered from nasal swabs was inconsistent; however, only 1/6 vaccinated NHPs (1572) had recoverable infectious virus on 3dpi, with no detectable infectious virus on 5 or 7dpi, while 3/6 control NHPs had positive TCID_50_ results at 3dpi, with 1/6 still having recoverable virus at 7dpi (Figure 5B). Mean viral RNA levels in rectal swabs increased from 1 to 3dpi in unvaccinated animals to approximately 10^3^ GEQ/mL and remained at that level until 7dpi. In contrast, only a single vaccinated animal, 1572, had positive rectal swabs on 1 and 3dpi, with all other vaccinated animals consistently being below the level of detection (Figure 5C). Infectious virus in rectal swabs was only recovered from a single, unvaccinated animal on day 7 (Figure 5D).

**Figure 5:**
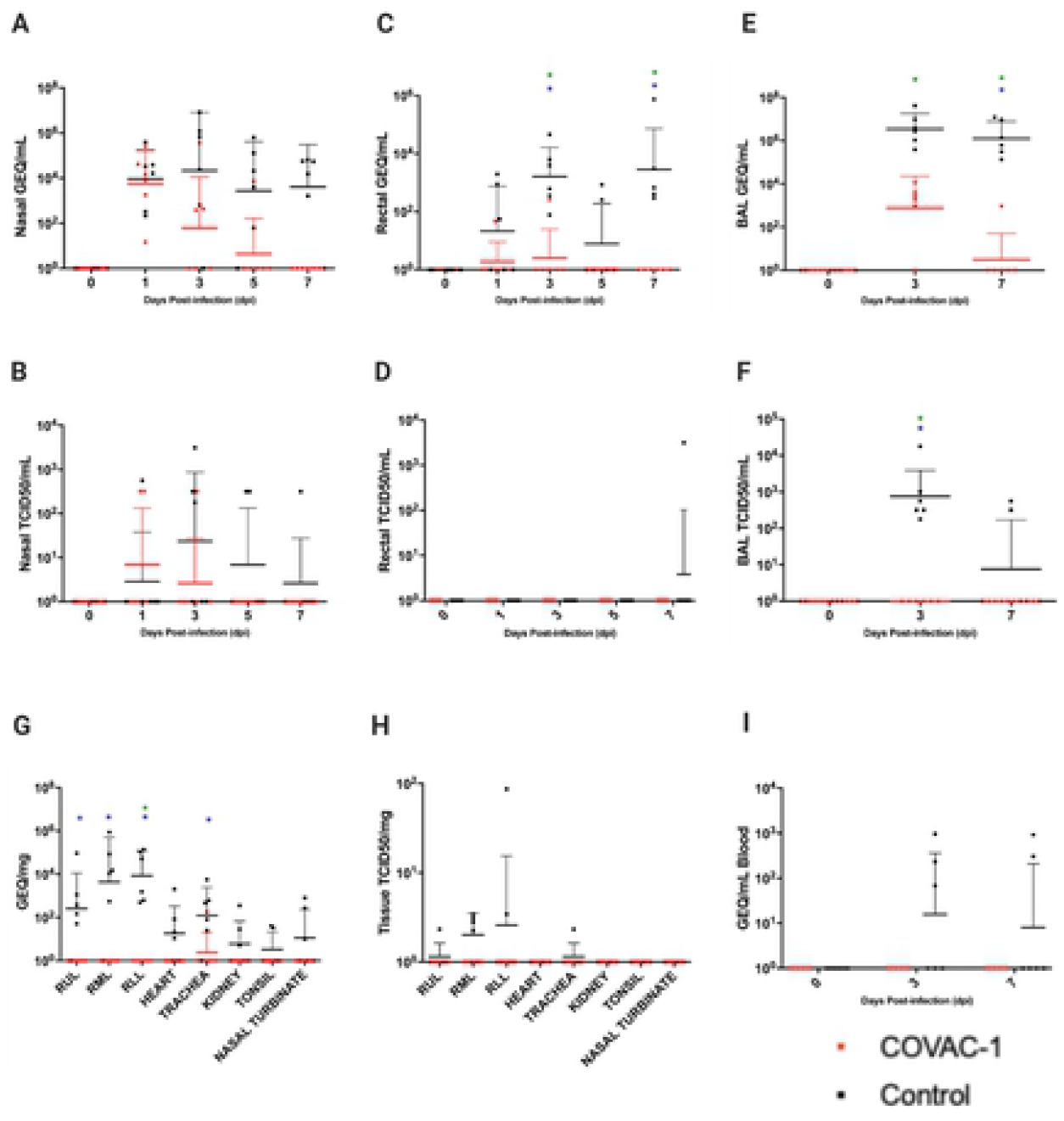
SARS-CoV-2 viral loads in control and COVAC-1 vaccinated AGMs. SARS-CoV-2 RNA and infectious virus in nasal swabs (A, B), rectal swabs (C, D), BAL fluid (E, F), tissues (G, H) and blood (I). Blue asterisk (*) indicates statistical significance (p<0.05) in percent positive when comparing COVAC-1 treated animals to unvaccinated controls. Green asterisk (*) represents a statistically significant difference in unequal means when comparing COVAC-1 to unvaccinated controls.

BAL fluid was also tested for the presence of viral RNA and infectious virus at 0, 3, and 7dpi. All control animals had high levels of viral RNA at 3dpi that were maintained until 7dpi. At 3dpi, the BAL samples from 5/6 control animals were positive for infectious virus, with 2 control animals still having detectable infectious virus on 7dpi (Figure 5E-F). In contrast, COVAC-1 vaccinated animals showed a significant reduction in mean viral RNA levels at 3dpi (p=0.00081) and 7dpi (p=0.000015), where only a single animal (1820) was found positive (Figure 5E). No infectious virus was recovered from the BAL of vaccinated animals at any time point, while 6/6 of unvaccinated animals were positive on 3dpi (p=0.0022), and 2/6 were still positive at 7dpi (Figure 5F). Very low levels of viral RNA were detected in the blood of 3/6 and 2/6 unvaccinated animals at 3 and 7 dpi, respectively (Figure 5H). Viral RNA was not detected in the blood of any of the vaccinated animals (Figure 5I).

Following euthanasia, collected tissues were evaluated for both viral RNA and infectious virus. Of the vaccinated group, only a single tissue sample collected from the trachea of animal 1512 was positive for viral RNA, with a low level of 1.83e+02 genome copies/mg (Figure 5G). All other tissues collected from vaccinated animals were negative. In contrast, all animals in the unvaccinated controls group had significantly higher levels of viral RNA in the sampled tissues, which included the right upper lung (RUL) (p=0.0152), right middle lung (RML) (p=0.0152), right lower lung (RLL) (p=0.0022) as well as nasal turbinate, trachea (p=0.0801), tonsil, heart and kidney, with five of six animals (1687, 1776, 1774, 1342, and 1512) having at least one sample positive for infectious virus as measured by TCID_50_ (Figure 5H).

### COVAC-1 prevents gross and histological pathological changes

Lung sections and heart, trachea, kidney, nasal turbinate, and tonsils were collected at 7dpi during necropsy. Gross pathology of the lungs of control animals consistently showed patchy areas of congestion, edema, and diffuse consolidation with areas of discoloration (Figure 6G). Alternatively, the control lungs were red in appearance and failed to collapse, indicative of inflammation. Comparable lung lesions were essentially absent in the vaccinated animals, where lungs appeared pink and readily collapsed (Figure 6H). No apparent changes were noted in other organs during gross pathology investigations.

**Figure 6:**
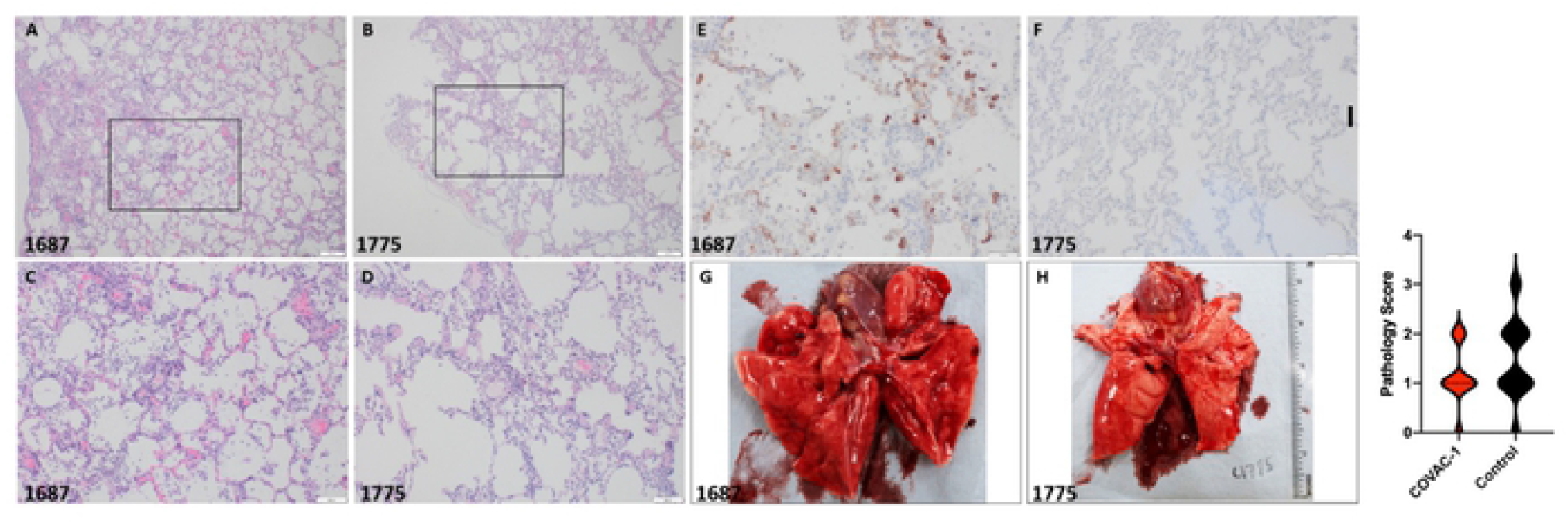
Representative histological staining and gross pathology of the right lower lung from control animal 1687 and COVAC-1 treated animal 1775. H&E staining at 4X magnification (A, B) and 10X magnification (C, D). Immunohistochemistry of the right lower lung of control animal 1687 showed strong immunoreactivity in pneumocytes and alveolar macrophages at 20X magnification (E). COVAC-1 treated animal 1775 showed an absence of staining in pneumocytes at 20X magnification (F). Gross pathology of control animal 1687 showed patchy red lesions indicative of inflammation and edema (G), which were not apparent in COVAC-1 treated animal 1775 (H). Histology additive pathology scores comparing COVAC-1 vaccinated animals and control animals (I).

Inflammation, type II pneumocyte hyperplasia and hemorrhage in the lungs were the most prominent features in animals infected with SARS-CoV-2. Histology was evaluated by additive pathology scores. Lung pathology was characterized by the presence of neutrophils and macrophages in bronchi/bronchioles, alveoli, and the interstitium. Some degree of this was observed in all animals. Both control and vaccinated animals had evidence of tracheal inflammation as well as neutrophils and macrophages in the bronchi, alveoli and/or interstitium in varying quantities. However, the control group had higher overall pathology scores when compared to the vaccinated animals, with the majority of samples (77%) from vaccinated animals scoring 0 or 1, with no scores of 3. In contrast, nearly half (47%) of samples from control animals scored above 1, with a small number (12%) scoring 3 (Figure 6I).

Lung sections from control animals collected on 7dpi were frequently positive for viral antigen staining (18/25 sections from the right middle and right lower lung (Figure 6E). One animal (1774) in the control group consistently had atypical staining on multiple occasions and was thus excluded from this analysis. Staining was observed in type II pneumocytes and less frequently in alveolar macrophages. In contrast, no positive antigen staining was observed in the same number of tissue sections in COVAC-1 vaccinated animals (Figure 6F).

### Serological responses

The serum IgG antibody response against SARS-CoV-2 was assessed using two antigens, soluble trimer or RBD by enzyme-linked immunosorbent assay (ELISA). All vaccinated animals had detectable IgG antibody responses in the soluble trimer ELISA, exceeding the upper limit of the assay (1:6400) 14 days prior to the challenge. Two vaccinated animals (1775, 1540) had decreased titres post-challenge, which subsequently increased by the end of the study. In contrast, all unvaccinated controls were negative for detectable IgG against soluble trimer. Five unvaccinated controls were also negative for detectable IgG against RBD throughout the study. However, one control animal (1774) had a titer of 1:400 following challenge on 7dpi. Vaccinated animals all reached the upper limit of 1:6400 42 days after vaccination (−14dpi) against RBD; however, animals 1775, 1820, 1478 and 1512 decreased slightly to 1:1600. By the end of the study, all animals except one (1775) had titres at the upper limit (1:6400) again (Table 1).

**Table 1:** IgG antibody responses following COVAC-1 vaccination and boost or PBS control and after SARS-CoV-2 challenge. Serum from animals at −14/-15 days pre-challenge and at 7dpi was assessed for binding to SARS-CoV-2 soluble trimer or RBD by IgG-specific ELISA.

| Animal (Treatment) | Soluble trimer<br>-14dpi/-15dpi | Soluble trimer<br>7dpi | RBD<br>-14dpi/-15dpi | RBD<br>7dpi |
| --- | --- | --- | --- | --- |
| 1164 (Control) | Negative | Negative | Negative | Negative |
| 1250 (Control) | Negative | Negative | Negative | Negative |
| 1342 (Control) | Negative | Negative | Negative | Negative |
| 1687 (Control) | Negative | Negative | Negative | Negative |
| 1774 (Control) | Negative | Negative | Negative | 1:400 |
| 1776 (Control) | Negative | Negative | Negative | Negative |
| 1478 (COVAC-1) | 1:6400 | 1:6400 | 1:6400 | 1:6400 |
| 1512 (COVAC-1) | 1:6400 | 1:6400 | 1:6400 | 1:6400 |
| 1540 (COVAC-1) | 1:6400 | 1:6400 | 1:6400 | 1:6400 |
| 1572 (COVAC-1) | 1:6400 | 1:6400 | 1:6400 | 1:6400 |
| 1775 (COVAC-1) | 1:6400 | 1:1600 | 1:6400 | 1:1600 |
| 1820 (COVAC-1) | 1:6400 | 1:6400 | 1:6400 | 1:6400 |

Neutralizing antibodies were assessed using a plaque reduction neutralization PRNT_50_ assay on serum from 14 days prior to virus challenge and 7 days following challenge, 42 and 63 days post-vaccination, respectively (Table 2). All six unvaccinated control animals were negative for neutralizing antibodies against SARS-CoV-2 (Canada/ON-VIDO-01/2020) 14 days before the virus challenge; however, by 7dpi, one control animal (1250) had detectable neutralizing antibody levels of 1:40. In contrast, all vaccinated animals had detectable neutralizing antibodies against SARS-CoV-2 (Canada/ON-VIDO-01/2020) at −14dpi with levels ranging from 1:20 to 1:160, with similar titres detected at 7dpi.

**Table 2:**
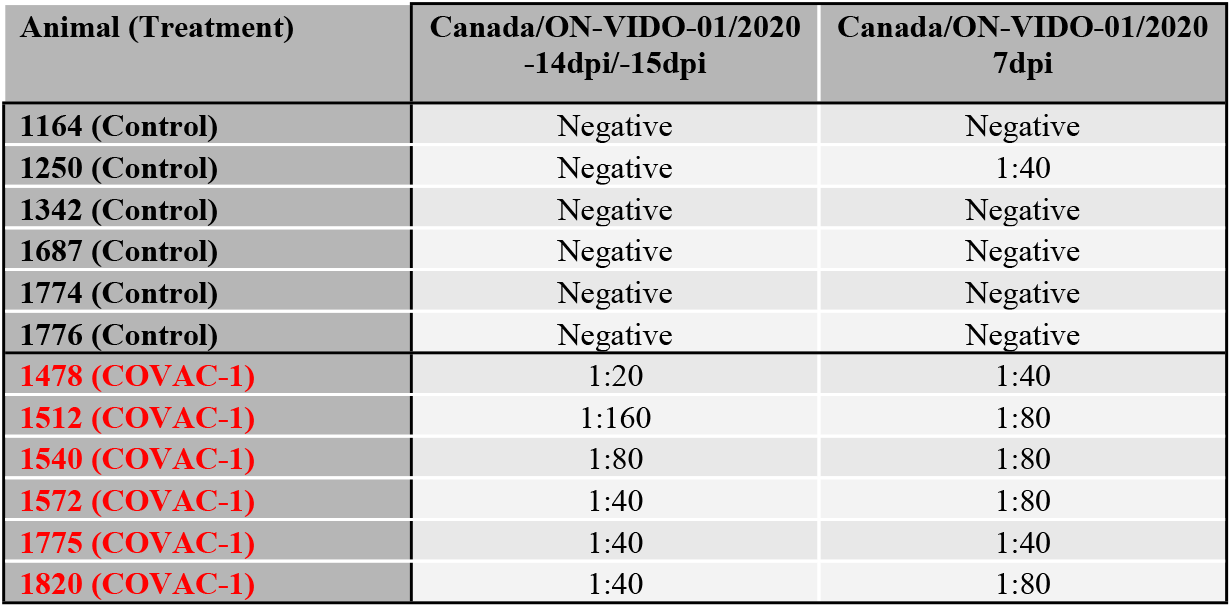
Neutralization 50% (PRNT50) assay using ancestral strain SARS-CoV-2

The ability of COVAC-1 to induce antibody-dependent cell-mediated cytotoxicity (ADCC) antibodies was also assessed. This was investigated as ADCC has been proposed to be a correlate of protection as it has been observed in individuals who appear to be protected from SARS-CoV-2 disease in the absence of neutralizing antibodies [25-27]. Low levels of antibodies capable of mediating ADCC were observed in 6/6 COVAC-1 treated animals 14 days following the boost vaccination (Figure 7A-B). The level of ADCC-mediating antibodies was maintained until the challenge and did not meaningfully change during the challenge period. In contrast, sera from control animals did not mediate ADCC activity at any tested time points, including following challenge. The breadth of antigen binding to the spike glycoproteins from other coronaviruses was also assessed separately. It was observed that sera from COVAC-1 vaccinated animals only bound to cells expressing SARS-CoV-2 S and that, similarly, binding was observed 14 days following the boost (on day 42) (Figure 7C-D). A much lower level of binding was also observed to SARS-CoV-1 but not to the spike glycoproteins from more distantly related coronaviruses, such as OC43, HKU1 or MERS-CoV (not shown).

**Figure 7:**
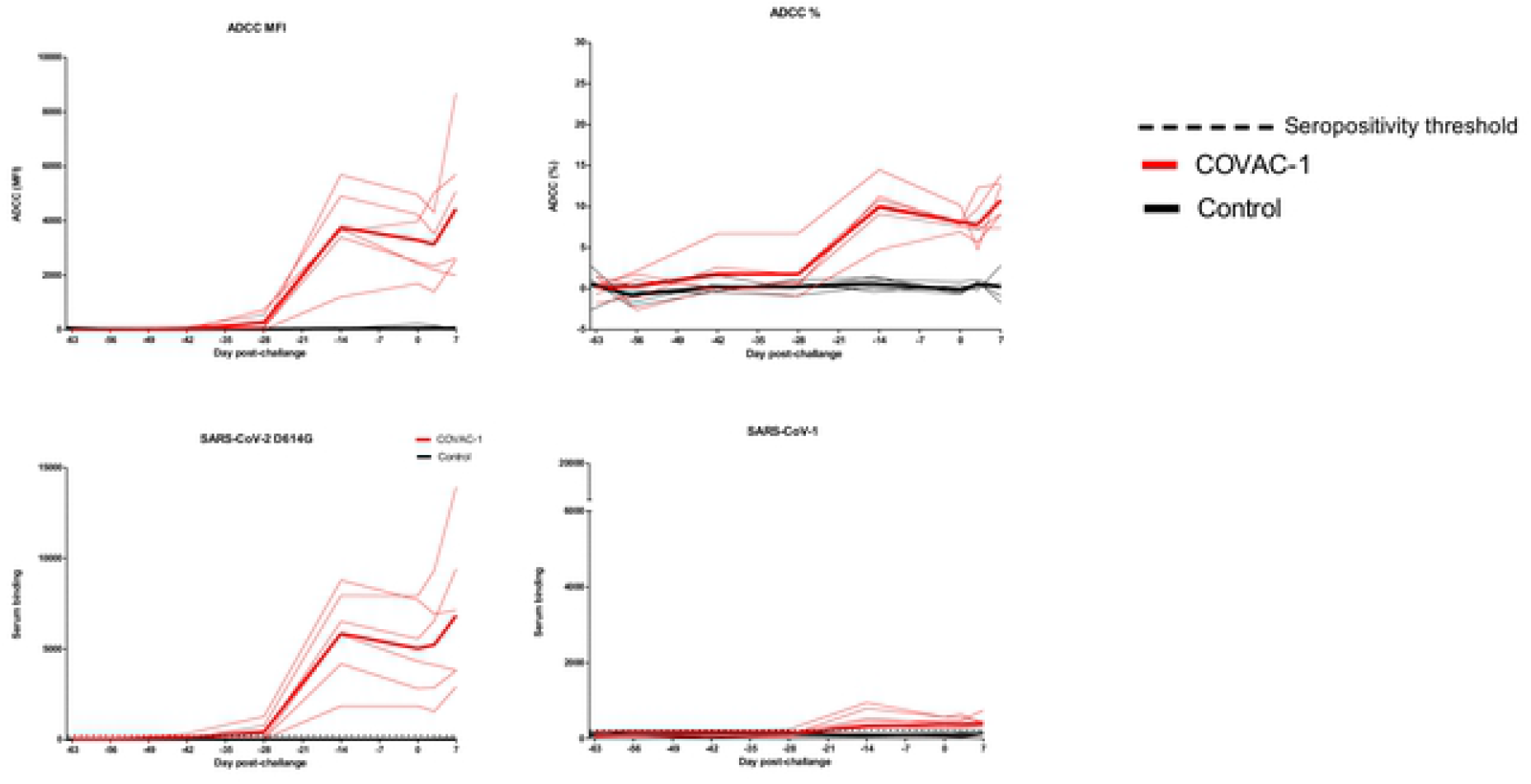
FACS-based ADCC assay was completed on CEM.NKr CCR5+ and CEM.NKr.SARS-CoV-2S target cells using serum from COVAC-1 vaccinated animals and unvaccinated control AGMs as effector cells. Results are presented as mean fluorescent intensity (MFI) (A), and percentage of ADCC obtained (B). Spike-specific binding was evaluated by flow cytometry using 293T cells expressing either SARS-CoV-2 (C) or SARS-CoV-1 (D) S full-length glycoproteins.

### Immune cell infiltration

Following the SARS-CoV-2 challenge, the extent of immune cell infiltration into BAL fluid (BALF) and blood was assessed by flow cytometry. Overall, the data supports that infiltration into the BALF occurred largely in the first few days following the challenge and substantially decreased thereafter.

In the COVAC-1 vaccinated group, HLA-DR+ CD8 T cells increased at a rate of 20% (95% CI: − 4 to 49%) per day, and although infiltration in the control group occurred at a significantly higher rate (p = 0.0495) of 66% (95% CI: 34 to 106%), none of the time points showed statistical significance when comparing the vaccinated and control groups (Figure 8A). COVAC-1 vaccinated animals had an increase in CD69+ CD8 T cells (34% per day, 95% CI: 9 to 65%; Figure 8B), however the unvaccinated controls also showed an increased rate (63% per day, 95% CI: 33 to 101%), resulting in no significant differences in slope (p = 0.2002). Despite this, the final concentration of CD69+ CD8 T cells at 7dpi was significantly different between the two groups, with the control group being approximately 9.7-fold higher (95% CI: 3 to 31-fold).

**Figure 8:**
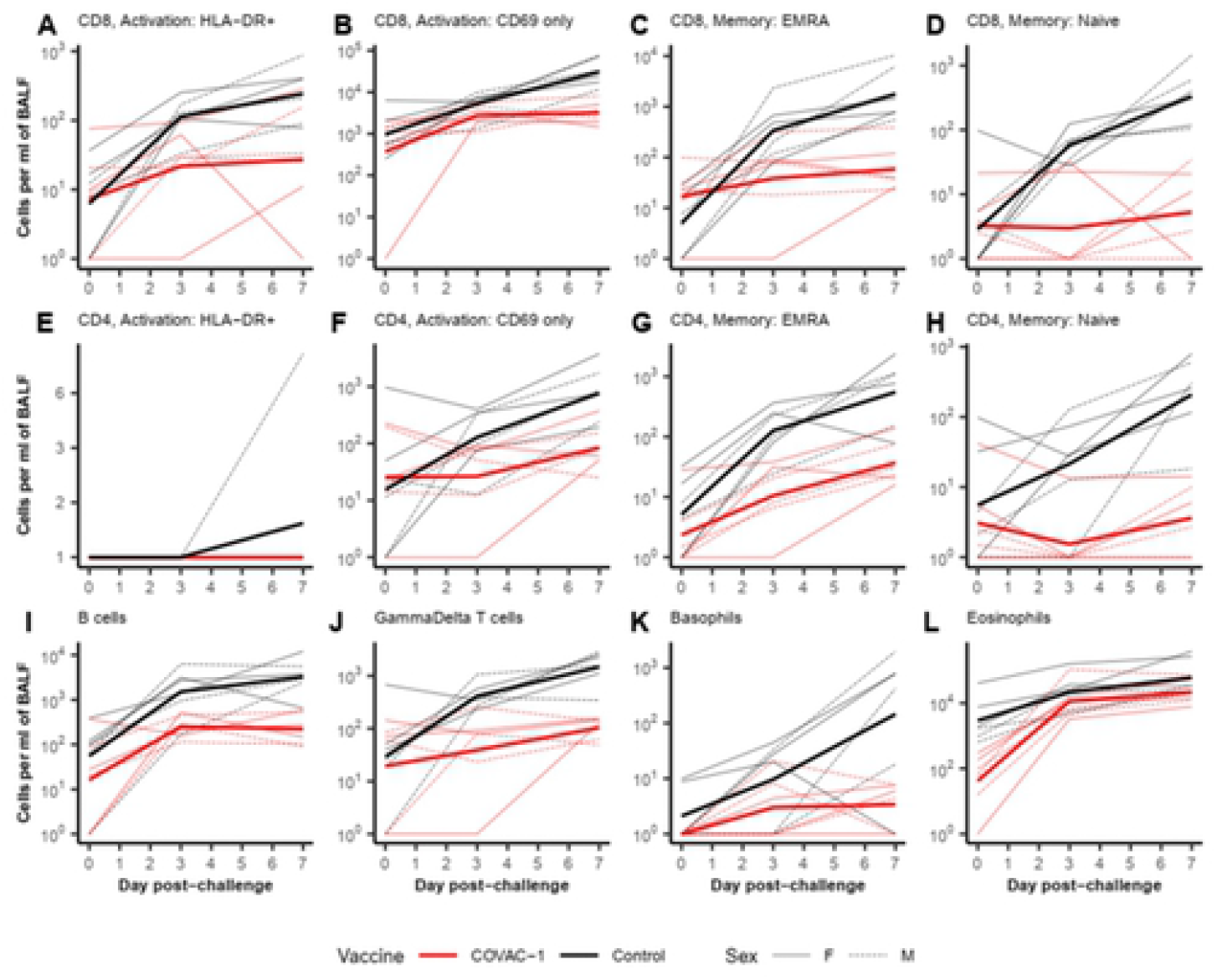
Infiltration of populations of interest in the BAL fluid of challenged animals. BAL fluid was processed by flow cytometry to assess the concentration of several cell populations. The plots present the concentration per mL of BAL fluid (+ 1 to account for 0s). Thin lines are individual animals, and thick lines are group averages. All other cell types evaluated are plotted and analyzed in Supplementary Material.

Interesting trends for the infiltration of memory phenotypes were also noted for both the terminally differentiated RA+ effector memory (EMRA) cells (Figure 8C) and the naïve cells (Figure 8D). In vaccinated animals, both cell populations show a similar pattern with little to no infiltration, a 19% increase per day for EMRA (95% CI: −2.7 to 46%); and a 7.7% increase per day for naïve cells (95% CI: −13.3 to 34%). However, the concentration of these cells in the BALF of the control group increased by 126% per day for EMRA (95% CI: 85 to 177%); and 95% increase per day for naïve cells (95% CI: 57 to 142%).

For CD4 T cells, no HLA-DR expression could be detected except in a single sample (Figure 8E). The COVAC-1 group had a negligible daily increase in CD69+ CD4 T cells (19% per day, 95% CI: −3.2 to 47%; Figure 8F), but the control group showed a significantly higher rate of infiltration (74% per day, 95% CI: 41 to 114%). The general trends for the EMRA (Figure 8G) and naïve (Figure 8H) CD4 T cells largely parallel those of the CD8 T cells. One distinction was that EMRA CD4 T cells infiltrated into the BALF of the COVAC-1 group at a significant rate (47% per day, 95% CI: 23 to 76%), while the infiltration in the control group was not significantly higher (92% per day, 95% CI: 61 to 129%). Despite this, the concentration of EMRA CD4 T cells was higher in the control group on both 3dpi (12-fold, 95% CI: 1.6 to 88) and 7dpi (15-fold higher, 95% CI: 2.3 to 99). As with the CD8 T cells, naïve CD4 T cells did not significantly infiltrate into the BALF in the COVAC-1 group (3% increase per day, 95% CI: −13 to 23%), but the infiltration into the BALF of the control group was significant (68% per day, 95% CI: 42 to 100%).

B cells infiltrated into the BALF of the COVAC-1 group at a rate of 42% per day (95% CI: 11 to 83%), but the infiltration in the BALF of the control group was not significantly higher, at 75% per day (95% CI: 36 to 125%) (Figure 8I). Similarly, γδ-T cells (Figure 8J), which play important roles in mucosal immunity, infiltrated into the BALF of the COVAC-1 group at a significant rate (27% per day, 95% CI: 1.7 to 59%). Still, the control group did not show significantly higher infiltration (73% per day, 95% CI: 38 to 116%). However, the control group still had significantly more γδ-T cells in its BALF (14-fold higher, 95% CI: 4.6 to 44) at 7dpi. Basophils (Figure 8K) did not infiltrate the BALF of the COVAC-1 group at a significant rate (18% per day, 95% CI: −7.3 to 51%) but did infiltrate the BALF of the control group at a significant rate (84% per day, 95% CI: 44 to 134%). Eosinophils showed a pattern opposite most of the other cell types; the infiltration was significantly higher in the COVAC-1 group (135% per day, 95% CI: 84 to 201%) than in the control group (52% per day, 95% CI: 19 to 95%) (Figure 8L); however, this may have been due to the control group having a significantly higher concentration of eosinophils on day 0 (30-fold higher, 95% CI: 25 to 36).

## DISCUSSION

Since the onset of the SARS-CoV-2 pandemic, accelerated trials have facilitated the approval of several vaccines; however, the pandemic remains an ongoing threat with a need for additional vaccines. Here we aimed to evaluate the immunogenicity and protective efficacy of a new S1 subunit vaccine and combination adjuvant, COVAC-1, in an AGM model of COVID-19. AGMs have been previously established as a disease model for SARS-CoV-2 to explore the dynamics of disease pathogenesis while recapitulating the human disease [15-17, 28]. Similar to previous studies, we observed that AGMs do not develop notable clinical illness, likely similar to many human cases. However, during gross and histological tissue pathology evaluation, they show pronounced damage of varying severity in respiratory tissues [15, 16]. In no case did the animals appear to have any form of distress, fever, weight loss, shivering, or any obvious discomfort. However, an interesting clinical finding that has not previously been documented was an increased pCO_2_ coinciding with an increased respiratory rate and no associated decrease in oxygen saturation that occurred in all the infected animals but was significantly higher in the control animals (Figure 2). In this context, this likely represents increased physiological dead space that may have manifested from the interstitial and airspace disease associated with the observed pathologic changes.

Also notable was that despite pathology being consistently observed in lung tissues of infected control animals, thoracic radiographic images taken throughout infection showed no evidence of pathological changes. Unremarkable X-rays, including during peak viremia, have been reported previously for SARS-CoV-2 infection in animal models and human patients [15, 29-31]. The alterations observed by X-ray may be more prominent following the early stages of the disease characterized by infection with SARS-CoV-2 when there are subsequently high levels of inflammation, coagulopathy and fibrosis [32]. Likely, clinical signs of infection, including crepitation and/or rales via auscultation (along with the corresponding pathologic correlates), may precede these radiographic signs. Unfortunately, the high containment level (BSL4) restrictions in this study design precluded auscultation but may be possible for future iterations.

Although clinical signs were absent to subtle in both vaccinated and control animals, thus not being useful as a comparator, viral indices (including RNA levels and TCID_50_) were significantly improved in vaccinated AGMs, suggesting protective efficacy from COVAC-1 vaccination. This protection was evident in the significantly reduced viral loads found in the BAL, mucosal swabs and tissues, an indication of reduced shedding and, therefore, potentially transmissibility as a result of vaccination [33]. While both groups had equivalent levels of viral RNA at 1dpi in nasal swabs, likely a result of the relatively high level of virus in the inoculum, all subsequent samples supported a rapid and significant reduction in viral RNA and infectious virus. This also included the absence of viral RNA in rectal swabs. Whether vaccination provides protection against virus replication in the gastrointestinal tract or whether the lack of viral RNA in rectal swabs in vaccinated animals is a result of overall less virus being produced in the upper respiratory and subsequently transiting the GI tract, both mechanisms support the notion that a good level of protection was achieved. Together, this supports that COVAC-1 may be effective at preventing or reducing transmission by reducing the amount of virus shed in the upper respiratory tract and excreted in feces.

This study’s histopathology of respiratory tissues showed varying degrees of inflammatory cell infiltration, including neutrophils and macrophages in bronchi/bronchioles, alveoli, and the interstitium, as observed previously in this model [16]. The trachea of multiple control AGMs and some of the vaccinated animals also showed mild to moderate levels of inflammation. However, tracheal inflammation may result from repeated BAL procedures throughout the study rather than from the infection itself. Notably, vaccination decreased the extent of pathological alterations in lung tissues compared to controls. Not only did this include lower overall scores, but also the absence of scores over 2. Consistent with decreased pathology and the absence of viral RNA, as assessed by RT-qPCR, was a complete absence of viral antigen in lung sections compared to consistently positive staining in control animals. Overall, this correlated with the gross pathological finding where vaccinated animals had essentially normal lungs in contrast to controls which showed congestion, edema, and diffuse consolidation with areas of discoloration.

Characterization of the cells that infiltrated into the lungs and were collected in BAL samples suggests that infiltration occurs early during infection. Similar processes occur in vaccinated and unvaccinated animals, with two notable differences. While not reaching significance due to large animal-to-animal variation, there was a consistent trend that control animals had overall higher levels of almost all characterized immune cell types, supportive of an overall higher level of immune cell infiltration. This data is consistent with histological data that also finds more immune cell infiltration and accompanying pathological alterations in the lungs of control animals. Perhaps expectedly, the second notable observation was that both naïve CD4 and CD8 cells were largely absent in vaccinated animals, while a memory response was active in vaccinated animals. While EMRA T cells also infiltrated the BAL fluid in control animals, this may be due to a non-specific response, given the large number of such cells involved, or cross-reactivity from previous coronavirus exposure [34, 35].

An important finding of this study is the production of ADCC-mediating antibodies in COVAC-1 vaccinated animals since the engagement of immune effector cells via non-neutralizing antibody functions, such as ADCC, is proposed to play an important role in the clearance of infected cells and protection from numerous viral pathogens [36-41]. In SARS-CoV-2, protection elicited following a single vaccination was associated with antibodies able to mediate ADCC [27]. Additionally, many infected human patients with mild disease have very low levels of neutralizing antibodies. In patients with severe disease that survive, ADCC-mediating antibodies are present at higher levels than in patients that do not survive [25, 42] and there is an inverse associated between ADCC and mortality [42]. This observation has even been extended to show an association between ADCC and survival following treatment with convalescent plasma [43]. Given the rapid clearance of the virus and the decreased pathological changes in the lungs, this would suggest that the COVAC-1 vaccine is sufficient to induce protection in this model. Not unexpectedly, antibody binding to other distantly related human coronaviruses such as OC43, HKU1 and MERS-CoV was not observed (not shown). However, a low level of binding to the SARS-CoV-1 spike glycoprotein was observed, suggesting that some cross-reactivity is present among more closely related coronaviruses.

Subunit vaccines offer several advantages over other vaccine formats that make them desirable. \ They are a proven technology that overall has excellent safety profiles. In clinical trials of the different vaccine platforms for COVID-19, local and systemic reaction rates were significantly lower among protein subunit vaccines than in three other COVID-19 vaccine platforms [44]. Subunit vaccines also retain a high safety profile even with multiple boosts [44-46]. Production is also highly scalable, and a significant global manufacturing infrastructure exists. However, in SARS-CoV-2, some limitations must still be overcome. The most critical of these may be ensuring that the spike glycoprotein is present in a form that elicits neutralizing antibodies, as non-neutralizing antibodies may mediate ADE. However, this has yet to be observed in the context of a polyclonal response [47-50]. In this study, S1 formulated with TriAdj, which typically induces a balanced to Th1-biased immune response [18], was demonstrated to provide a high level of protection against the SARS-CoV-2 challenge. In addition, no adverse effects were noted in any of the vaccinated AGMs.

However, there are several limitations with this animal model and in this study. This study evaluated gross and histological pathology at limited terminal time points, perhaps missing aspects of disease pathogenesis in the control or vaccinated groups. In addition, the AGM model of SARS-CoV-2 does not show significant clinical disease. Thus, it is not possible to conclude that vaccination prevents severe disease, despite this being inferred given the rapid clearance of the virus. Moreover, it has been quite apparent from the human cases that severe disease manifests more frequently with associated comorbidities and at later times in the disease course that we may have precluded with early termination of this experiment. However, it was anticipated that comparing levels of virus in tissues would no longer be possible at later times.

In summary, we showed effective immunogenicity and protective effects against SARS-CoV-2 challenge from two injections of an S1 subunit vaccine candidate containing TriAdj (COVAC-1). Specifically, our evaluation demonstrated that COVAC-1 had no apparent side effects or ADEs, had a protective effect from lung disease pathology, and elicited neutralizing antibodies in sera of immunized AGMs against SARS-CoV-2 (Canada/ON-VIDO-01/2020). Moreover, vaccinated animals had significantly reduced viral loads within BAL, mucosal swabs, and tissues compared to control animals. This vaccine candidate should be considered for further investigation and development, contributing to the armamentarium in the worldwide fight against SARS-CoV-2.

## METHODOLOGY

### Vaccine

COVAC-1 consists of a codon-optimized mammalian-produced HIS-tagged S1 protein produced by Biodextris (400µg/mL Lot: C2003-VID-DSP-E-002) that has been formulated with TriAdj comprising 250µg PCEP, 250µg poly I:C (Lot: PJ625E01) and 500µg IDR-1002 per dose in PBS. Six African green monkeys (AGM) (3 female: 3 male) (animal numbers: 1478, 1512, 1540, 1572, 1775, 1820) received 50µg of mammalian-produced SARS-CoV-2 S1 formulated with TriAdj in a volume of 0.5mL delivered via the intramuscular (i.m.) route on the caudal thigh. Animals were vaccinated on day −56 and boosted on day −28 pre-challenge. Six control AGMs (3 female: 3 male) (animal numbers 1164, 1250, 1342, 1687, 1774, 1776) were mock vaccinated with PBS following the same sampling timeline as the vaccinated animals.

### Animal Challenge

All the animal experiments were approved by the Animal Care Committees of the Canadian Science Center for Human and Animal Health and the University of Saskatchewan in accordance with the guidelines provided by the Canadian Council on Animal Care. All experiments with live SARS-CoV-2 were completed within the Biosafety Level 3 (BSL3) or BSL4 laboratories. All twelve AGMs were inoculated with target dose of 5E+04 TCID_50_ per animal of SARS-CoV-2 (Canada/ON-VIDO-01/2020, GISAID #EPI_ISL_425177) delivered as follows: 0.5mL per nare intranasal (i.n), 5mL intratracheal (i.t.), 1mL oral and 0.2mL per eye ocular routes (i.o) for a total volume of 7.4mL. The inoculum was back titered on Vero cells and indicated a total challenge dose of 1.54E+04 TCID_50_ was delivered. Animals were monitored twice daily for clinical signs of illness, including fever, clinical appearance/behaviour, and respiratory signs. All procedures requiring handling were performed under sedation by ketamine or ketamine in addition to dexmedetomidine.

### *In vivo* sampling

Vaccinated animals were sedated, and blood was collected on days −56 (vaccination), −49, −42, −28 (vaccine boost), −14, 0, 3, and 7 days post-infection (dpi). Oral, rectal, and nasal swabs (MedPro 018-430) were taken to assess viral shedding at 0, 1, 3, 5, and 7 dpi. Bronchoalveolar lavages (BAL) were completed at days −49, 0, 3 and 7 post-challenge. Electrocardiogram (ECG) and X-rays were performed on days 0, 1, 3, 5 and 7 post infection (Figure 1A). Unvaccinated controls followed a similar sampling timeline; however, the viral challenge with SARS-CoV-2 was delayed by one day. Therefore, sampling days included −57, − 50, −43, −29, −15 before challenge and 0, 1, 3, 5 and 7 post virus challenge (Figure 1A).

BAL procedures were completed on ketamine and dexmedetomidine-sedated animals receiving 100% O_2_. Animals were intubated with a 3.5-4.5mm cuffed endotracheal tube (ETT) (COVIDIEN, 86445). Subsequently, a sterile suction catheter was placed past the end of an inserted ETT into a main stem bronchus in the distal airway. Saline was then infused through the catheter and immediately aspirated back into the syringe. Once the initial fluid was recovered, the process was repeated 2-3 times, followed by the removal of the catheter. Adequate yields for this procedure are ∼60% of the total infused volume.

### Virus inactivation, RNA extraction, RT-qPCR

140 µl of fluid samples were inactivated with AVL (QIAGEN, Valencia, CA), and RNA was subsequently extracted using the QIAmp viral RNA Mini kit (QIAGEN, Valencia, CA) following the manufacturer’s instructions. For tissue samples, 30mg of homogenized tissue was inactivated with RLT and extracted using the RNeasy Mini kit (QIAGEN, Valencia, CA) following manufacturer instructions. Primers and probes used to detect SARS-CoV-2 by real-time quantitative PCR (RT-qPCR) were based on the E gene (E_Sarbeco_F1: ACAGGTACGTTAATAGTTAATAGCGT, E_Sarbeco_R2: ATATTGCAGCAGTACGCACACA, E_Sarbeco_P1: FAM-ACACTAGCCATCCTTACTGCGCTTCG-BHQ), described by Corman et al. [51]. RT-qPCR was performed using TaqPath master mix (ThermoFisher Scientific) and run on a QuantStudio 5 RT-qPCR system to measure the cycle threshold (Ct).

To compare differences in viral RNA levels between the COVAC-1 vaccinated group and the control group, a Fisher’s exact test was completed using GraphPad Prism 9. Additionally, an unpaired t-test with Welch correction was used to analyze the difference in unequal means. The outcome variable was transformed as log_10_(GEQ/mL + 1) to account for samples with 0 viral genome copies.

### Virus Titration

Virus titration was performed by tissue culture infectious dose 50 (TCID_50_) assay using Vero cells (ATCC CCL-81) on blood, swab, BAL and tissue samples that had a Ct at or below 27 measured by RT-qPCR [33]. Briefly, increasing 10-fold dilutions of the samples were incubated on Vero monolayers maintained in modified Eagle’s medium (MEM) supplemented with 5% fetal bovine serum (FBS), 1% penicillin/streptomycin, 1% L-glutamine, in triplicate and incubated at 37°C with 5% CO_2_. Following incubation for 96-120 hours, the cytopathic effect was measured under a microscope, and TCID_50_/mL or mg was calculated using the Reed and Muench method as previously described [52].

To compare differences in infectious virus levels between the COVAC-1 vaccinated group and the control group, a Fisher’s exact test was completed using GraphPad Prism 9. Furthermore, an unpaired t-test with Welch correction was used to analyze the difference in unequal means. The outcome variable was transformed as log_10_(TCID_50_/mL + 1) to account for samples with 0 infectious virus recovered.

### Hematology

Whole blood collected in an EDTA tube (BD vacutainer) was used to test total white blood cell (WBC) counts, cell differential distribution, red blood cell counts, platelet counts, hematocrit values, total hemoglobin concentrations and other blood markers using an HM5 hematologic analyzer (Abaxis). Additionally, lithium heparin treated blood was tested for markers of organ function, specifically liver and kidney, including albumin (ALB), amylase, alanine aminotransferase (ALT), alkaline phosphatase (ALP), blood urea nitrogen (BUN), calcium, creatinine (CRE), using a Vetscan (Abaxis) and a piccolo point-of-care analyzer with Metlac and Biochemistry Panel Plus analyzer discs (Abaxis). Blood gas analysis, including partial pressures of CO_2_ and O_2_, pH, bicarbonate, glucose, sodium and potassium levels, was obtained using an iSTAT Alinity hematological analyzer (Abbott).

### ELISA

Serum collected at the time points indicated above were tested for SARS-CoV-2 specific antibodies against soluble trimer (S1+S2) (Sino Biological 40859-V08H1) and RBD (Sino Biological 40592-V08B) recombinant proteins. HI BIND Assay plates (COSTAR 3366) were coated with antigen diluted 1:1000 in PBS overnight at 4°C. Plates were then washed three times with wash buffer (PBS with 0.1% Tween20). Wells were then blocked with 100µL of 5% skim milk in 0.2% Tween20 (DIFCO BD 232100) in PBS at 37°C. After blocking, serum samples were four-fold serially diluted from 1:100 down to 1:6400 in 5% skim milk in 0.2% Tween20 solution in the pre-coated wells and incubated at 37°C for 1 hour. Serum collected at days −56 and −57 post infection for the vaccinated and control animal was used as a baseline. Plates were subsequently washed with wash buffer three times, and then 100µL of HRP labelled goat anti-human IgG (SERA CARE 5450-0009), diluted 1:2000 in 5% skim milk in 0.2% Tween20 in PBS, was added to the wells and placed at 37°C for 1 hour followed by three more washes. After the final wash, KPL ABST substrate solution A (SERA CARE 5210-0035) and peroxidase substrate solution B (SERA CARE 5120-0038) were mixed in a 1:1 ratio, and 100µL was added to each well. The plates were then incubated for 30 minutes at 37°C, and absorbance values were determined at 405nm.

### Serum Neutralization Assay

Neutralization titres were calculated by determining the dilution of serum that reduced plaques by 50% (PRNT_50_) following the previously outlined methodology [53]. Briefly, serum collected at the sampling times outlined above was diluted 2-fold and incubated with SARS-CoV-2 (Canada/ON-VIDO-01/2020) for 1 hour at 37°C with 5% CO_2_. 100µL of the inoculum was then placed on Vero E6 cells (ATCC CRL-1586) for 1 hour at 37°C with 5% CO_2_, rocking every 15 minutes. The cells were then overlaid with a 3% carboxymethylcellulose (CMC) liquid matrix diluted 1:1 with 2X MEM supplemented with 8% FBS, 2% L-glutamine, 2% penicillin and streptomycin, 2% non-essential amino acids, 7.5% sodium bicarbonate and incubated at 37°C for 72 hours. Following incubation, the CMC was aspirated, and the cell monolayer was fixed for 1 hour with 10% buffered formalin. The formalin was then removed, and 100µL of 0.5% crystal violet (CV) solution was added to each well for 15 minutes to stain the cells. 1mL of 20% ethanol was added to each well for washing, followed by aspiration of both the CV and ethanol before plaques were counted.

### ADCC Assay

A detailed STAR Protocol is available for the ADCC assay [54]. Briefly, parental CEM.NKr CCR5+ cells were mixed at a 1:1 ratio with CEM.NKr.SARS-CoV-2.S cells and stained for viability with AquaVivid (Thermo Fisher Scientific, Waltham, MA, USA) and with cell proliferation dye eFluor670 (Thermo Fisher Scientific) to generate target cells. Overnight rested human PBMCs were stained with cell proliferation dye eFluor450 (Thermo Fisher Scientific) and used as effector cells. Stained target and effector cells were mixed at a ratio of 1:10 in 96-well V-bottom plates. Gamma irradiated serum from vaccinated and control AGMs (1/500 dilution) were added to the appropriate wells. The plates were subsequently centrifuged for 1 min at 300 g and incubated at 37°C, 5% CO_2_ for 5 h prior to being fixed with 2% PBS-formaldehyde. Samples were acquired on an LSRII cytometer (BD Biosciences), and data analysis was performed using FlowJo v10.7.1 (Tree Star). ADCC activity was calculated as previously described [54].

### Cell-based Spike Binding Assay

This assay was performed as previously described (34). Briefly, 293T cells were transfected individually with a plasmid encoding the indicated S glycoprotein (D614G, SARS-CoV-1, OC43, HKU1, MERS-CoV). 48 h post-transfection, S-expressing cells were stained with the CV3-25 Ab or gamma-irradiated serum from vaccinated or control AGMs (1/250 dilution) followed by AlexaFluor-647-conjugated goat anti-human IgM+IgG+IgA Abs (1/800 dilution) as secondary Abs. The percentage of transduced cells (GFP+ cells) was determined by gating the living cell population based on viability dye staining (Aqua Vivid, Invitrogen). Samples were acquired on an LSRII cytometer (BD Biosciences), and data analysis was performed using FlowJo v10.7.1 (Tree Star). The seropositivity threshold was established as previously described [27].

### Histopathology and Immunohistochemistry

Tissues collected during necropsy, approximately 1cm^3^ in size, were immersion-fixed in 10% neutral buffered formalin. Tissues collected included right upper, middle and lower lung, heart, trachea, kidney, tonsils and nasal turbinate. Lung tissues were submitted in tissue cassettes and processed overnight in a Sakura Tissue-Tek VIP6AI Tissue Processor. Samples were taken from 10% neutral buffered formalin, through increasing concentrations of alcohol, to xylene, then finished in paraffin wax over 14 hours. Samples were taken from the tissue cassettes and placed in metal moulds filled with molten wax to create a paraffin “block”. Blocks were subsequently cut at 4 µm on a microtome.

### Hematoxylin and Eosin

Hematoxylin and Eosin (H&E) staining was performed on a Sakura (Tissue-Tek) Prisma Automated Slide Stainer. Slides were placed on slide staining racks and deparaffined for 1 hour at 60°C. Surgipath Hematoxylin 560, Surgipath Blue Buffer 8 and Surgipath Alcohol Eosin Y 515 were used onboard the automated stainer. Slides were coverslipped using the Sakura (Tissue-Tek) Glas G2 Automated Coverslipper with Somagen Tissue-Tek Glas Mounting Medium. Tissue sections, including upper, middle, and lower right and left lungs, were scored based on inflammation (0: absent, 1: slight or questionable, 2: clearly present, 3: moderate, 4: severe), the proportion of parenchyma affected, the extent of hypertrophy of alveolar pneumocytes and intensity of hemorrhage.

### Immunohistochemistry

Lung tissue sections were prepared for immunohistochemical staining, conducted at Prairie Diagnostic Services (Saskatoon, SK) using an automated slide stainer (Autostainer Plus, Agilent Technologies Canada Inc., Mississauga, ON). Epitope retrieval was performed in Tris/EDTA, pH 9, at 97°C for 20 minutes. The primary antibody was a rabbit polyclonal antibody against the nucleocapsid protein of SARS-CoV-2 (SARS2-N). The SARS2-N-specific antibody was produced in-house by VIDO (Animal Study number AS#20-012). The SARS2-N-specific antibody was diluted at 1:800 in PBS and incubated with the slides for 30 minutes at room temperature. After washing, the bound SARS2-N antibody was then detected using an HRP-labelled polymer detection reagent (EnVision+ System - HRP Labelled Polymer, Agilent Technologies Canada Inc., Mississauga, ON). Immunostaining was categorized as no staining, weak staining intensity, or strong staining intensity.

### Flow cytometry and Immunophenotyping

The BALF was centrifuged at 600 x g for 10 minutes. The pellet was then resuspended in 0.5mL plain RPMI medium. An aliquot was mixed with an equal volume of PBS 5 µg/ml Acridine Orange (Life Technologies) 100 µg/mL Propidium Iodide (Life Technologies) and counted on a Nexcelom Cellometer Auto 2000. One hundred microliters of resuspended BALF cells or whole blood were stained as described in the “Immunophenotyping for NHPs, containment protocol” [55]. The samples were run on a FACSymphony A5 instrument.

The counts for each population of interest were normalized to the Live CD45+ cells and multiplied by the number of cells per mL of fluid to obtain the concentration of each cell type in the BALF or the count of white blood cells (WBC) from the HM5 analyzer. To compare the cell concentration between the vaccinated and control groups at each time point, a two-way repeated measures mixed-effect ANOVA with Šídák’s multiple comparisons test was performed using GraphPad Prism 9. The overall trend over time across groups was compared by using a linear mixed effect model in R. The outcome variable was the log_10_(Cell Concentration + 1) to account for samples with a cell concentration of 0.

## ACKNOWLEDGEMENTS

SARS-CoV-2 research is supported in the laboratory of D.F. by the Canadian Institutes of Health Research (CIHR) grants OV5-170349, VRI-173022 and VS1-175531 and in the laboratory of A.F by grants 352417 and 177958 and by an Exceptional Fund COVID-19 from the Canada Foundation for Innovation (CFI) #41027 to A.F. D.F. is a member of the CIHR-funded Coronavirus Variants Rapid Response Network (CoVaRR-Net). VIDO receives operational support from the Province of Saskatchewan through Innovation Saskatchewan and from the Canadian Foundation for Innovation through the Major Science Initiatives. A.F. is the recipient of a Canada Research Chair on Retroviral Entry # RCHS0235 950-232424. G.B.B. is the recipient of a Fonds de Recherche Québec-Santé (FRQS) PhD fellowship. The funders had no role in study design, data collection and analysis, decision to publish, or preparation of the manuscript. The authors have declared that no competing interests exist.

## REFERENCES

1. Wang Y, Wang L, Cao H, Liu C. SARS-CoV-2 S1 is superior to the RBD as a COVID-19 subunit vaccine antigen. J Med Virol. 2021;93(2):892-8. Epub 2020/07/22. doi: 10.1002/jmv.26320. PubMed PMID: 32691875; PubMed Central PMCID: PMCPMC7404424.

2. Knoll MD, Wonodi C. Oxford-AstraZeneca COVID-19 vaccine efficacy. Lancet. 2021;397(10269):72-4. Epub 2020/12/12. doi: 10.1016/S0140-6736(20)32623-4. PubMed PMID: 33306990; PubMed Central PMCID: PMCPMC7832220.

3. Polack FP, Thomas SJ, Kitchin N, Absalon J, Gurtman A, Lockhart S, et al. Safety and Efficacy of the BNT162b2 mRNA Covid-19 Vaccine. N Engl J Med. 2020;383(27):2603-15. Epub 2020/12/11. doi: 10.1056/NEJMoa2034577. PubMed PMID: 33301246; PubMed Central PMCID: PMCPMC7745181.

4. Al Kaabi N, Zhang Y, Xia S, Yang Y, Al Qahtani MM, Abdulrazzaq N, et al. Effect of 2 Inactivated SARS-CoV-2 Vaccines on Symptomatic COVID-19 Infection in Adults: A Randomized Clinical Trial. JAMA. 2021;326(1):35-45. Epub 2021/05/27. doi: 10.1001/jama.2021.8565. PubMed PMID: 34037666; PubMed Central PMCID: PMCPMC8156175.

5. Lazaro-Frias A, Perez P, Zamora C, Sanchez-Cordon PJ, Guzman M, Luczkowiak J, et al. Full efficacy and long-term immunogenicity induced by the SARS-CoV-2 vaccine candidate MVA-CoV2-S in mice. NPJ Vaccines. 2022;7(1):17. Epub 2022/02/11. doi: 10.1038/s41541-022-00440-w. PubMed PMID: 35140227; PubMed Central PMCID: PMCPMC8828760.

6. Escriou N, Callendret B, Lorin V, Combredet C, Marianneau P, Fevrier M, et al. Protection from SARS coronavirus conferred by live measles vaccine expressing the spike glycoprotein. Virology. 2014;452-453:32-41. Epub 2014/03/13. doi: 10.1016/j.virol.2014.01.002. PubMed PMID: 24606680; PubMed Central PMCID: PMCPMC7111909.

7. Kim YI, Kim D, Yu KM, Seo HD, Lee SA, Casel MAB, et al. Development of Spike Receptor-Binding Domain Nanoparticles as a Vaccine Candidate against SARS-CoV-2 Infection in Ferrets. mBio. 2021;12(2). Epub 2021/03/04. doi: 10.1128/mBio.00230-21. PubMed PMID: 33653891; PubMed Central PMCID: PMCPMC8092224.

8. Prompetchara E, Ketloy C, Tharakhet K, Kaewpang P, Buranapraditkun S, Techawiwattanaboon T, et al. DNA vaccine candidate encoding SARS-CoV-2 spike proteins elicited potent humoral and Th1 cell-mediated immune responses in mice. PLoS One. 2021;16(3):e0248007. Epub 2021/03/23. doi: 10.1371/journal.pone.0248007. PubMed PMID: 33750975; PubMed Central PMCID: PMCPMC7984610 following competing interests to declare: WW is a consultant for BioNet-Asia Co., Ltd. BioNet-Asia Co., Ltd provided peptides for T cells assay. This does not alter our adherence to PLOS ONE policies on sharing data and materials. There are no patents, products in development or marketed products associated with this research to declare.

9. Martinez-Flores D, Zepeda-Cervantes J, Cruz-Resendiz A, Aguirre-Sampieri S, Sampieri A, Vaca L. SARS-CoV-2 Vaccines Based on the Spike Glycoprotein and Implications of New Viral Variants. Front Immunol. 2021;12:701501. Epub 2021/07/30. doi: 10.3389/fimmu.2021.701501. PubMed PMID: 34322129; PubMed Central PMCID: PMCPMC8311925.

10. Kaur SP, Gupta V. COVID-19 Vaccine: A comprehensive status report. Virus Res. 2020;288:198114. Epub 2020/08/18. doi: 10.1016/j.virusres.2020.198114. PubMed PMID: 32800805; PubMed Central PMCID: PMCPMC7423510.

11. Guo Y, He W, Mou H, Zhang L, Chang J, Peng S, et al. An Engineered Receptor-Binding Domain Improves the Immunogenicity of Multivalent SARS-CoV-2 Vaccines. mBio. 2021;12(3). Epub 2021/05/13. doi: 10.1128/mBio.00930-21. PubMed PMID: 33975938; PubMed Central PMCID: PMCPMC8262850.

12. Liu Z, Xu W, Xia S, Gu C, Wang X, Wang Q, et al. RBD-Fc-based COVID-19 vaccine candidate induces highly potent SARS-CoV-2 neutralizing antibody response. Signal Transduct Target Ther. 2020;5(1):282. Epub 2020/11/29. doi: 10.1038/s41392-020-00402-5. PubMed PMID: 33247109; PubMed Central PMCID: PMCPMC7691975.

13. He Y, Zhou Y, Liu S, Kou Z, Li W, Farzan M, et al. Receptor-binding domain of SARS-CoV spike protein induces highly potent neutralizing antibodies: implication for developing subunit vaccine. Biochem Biophys Res Commun. 2004;324(2):773-81. Epub 2004/10/12. doi: 10.1016/j.bbrc.2004.09.106. PubMed PMID: 15474494; PubMed Central PMCID: PMCPMC7092904.

14. Reed SG, Orr MT, Fox CB. Key roles of adjuvants in modern vaccines. Nat Med. 2013;19(12):1597-608. Epub 2013/12/07. doi: 10.1038/nm.3409. PubMed PMID: 24309663.

15. Woolsey C, Borisevich V, Prasad AN, Agans KN, Deer DJ, Dobias NS, et al. Establishment of an African green monkey model for COVID-19 and protection against re-infection. Nat Immunol. 2021;22(1):86-98. Epub 2020/11/26. doi: 10.1038/s41590-020-00835-8. PubMed PMID: 33235385; PubMed Central PMCID: PMCPMC7790436.

16. Speranza E, Williamson BN, Feldmann F, Sturdevant GL, Perez-Perez L, Meade-White K, et al. Single-cell RNA sequencing reveals SARS-CoV-2 infection dynamics in lungs of African green monkeys. Sci Transl Med. 2021;13(578). Epub 2021/01/13. doi: 10.1126/scitranslmed.abe8146. PubMed PMID: 33431511; PubMed Central PMCID: PMCPMC7875333.

17. Cross RW, Agans KN, Prasad AN, Borisevich V, Woolsey C, Deer DJ, et al. Intranasal exposure of African green monkeys to SARS-CoV-2 results in acute phase pneumonia with shedding and lung injury still present in the early convalescence phase. Virol J. 2020;17(1):1-12. Epub 2020/08/21. doi: 10.21203/rs.3.rs-50023/v2. PubMed PMID: 32818211; PubMed Central PMCID: PMCPMC7430587.

18. Garg R, Latimer L, Gerdts V, Potter A, van Drunen Littel-van den Hurk S. Vaccination with the RSV fusion protein formulated with a combination adjuvant induces long-lasting protective immunity. J Gen Virol. 2014;95(Pt 5):1043-54. Epub 2014/02/28. doi: 10.1099/vir.0.062570-0. PubMed PMID: 24572813.

19. Garg R, Brownlie R, Latimer L, Gerdts V, Potter A, van Drunen Littel-van den Hurk S. A chimeric glycoprotein formulated with a combination adjuvant induces protective immunity against both human respiratory syncytial virus and parainfluenza virus type 3. Antiviral Res. 2018;158:78-87. Epub 2018/08/03. doi: 10.1016/j.antiviral.2018.07.021. PubMed PMID: 30071204.

20. Garg R, Latimer L, Gomis S, Gerdts V, Potter A, van Drunen Littel-van den Hurk S. Maternal vaccination with a novel chimeric glycoprotein formulated with a polymer-based adjuvant provides protection from human parainfluenza virus type 3 in newborn lambs. Antiviral Res. 2019;162:54-60. Epub 2018/12/15. doi: 10.1016/j.antiviral.2018.12.010. PubMed PMID: 30550799.

21. Wilson HL, Kovacs-Nolan J, Latimer L, Buchanan R, Gomis S, Babiuk L, et al. A novel triple adjuvant formulation promotes strong, Th1-biased immune responses and significant antigen retention at the site of injection. Vaccine. 2010;28(52):8288-99. Epub 2010/10/21. doi: 10.1016/j.vaccine.2010.10.006. PubMed PMID: 20959153.

22. Lu Y, Landreth S, Liu G, Brownlie R, Gaba A, Littel-van den Hurk SVD, et al. Innate immunemodulator containing adjuvant formulated HA based vaccine protects mice from lethal infection of highly pathogenic avian influenza H5N1 virus. Vaccine. 2020;38(10):2387-95. Epub 2020/02/06. doi: 10.1016/j.vaccine.2020.01.051. PubMed PMID: 32014270.

23. Garg R, Babiuk L, van Drunen Littel-van den Hurk S, Gerdts V. A novel combination adjuvant platform for human and animal vaccines. Vaccine. 2017;35(35 Pt A):4486-9. Epub 2017/06/11. doi: 10.1016/j.vaccine.2017.05.067. PubMed PMID: 28599794.

24. Blair RV, Vaccari M, Doyle-Meyers LA, Roy CJ, Russell-Lodrigue K, Fahlberg M, et al. Acute Respiratory Distress in Aged, SARS-CoV-2-Infected African Green Monkeys but Not Rhesus Macaques. Am J Pathol. 2021;191(2):274-82. Epub 2020/11/11. doi: 10.1016/j.ajpath.2020.10.016. PubMed PMID: 33171111; PubMed Central PMCID: PMCPMC7648506.

25. Yu Y, Wang M, Zhang X, Li S, Lu Q, Zeng H, et al. Antibody-dependent cellular cytotoxicity response to SARS-CoV-2 in COVID-19 patients. Signal Transduct Target Ther. 2021;6(1):346. Epub 2021/09/26. doi: 10.1038/s41392-021-00759-1. PubMed PMID: 34561414; PubMed Central PMCID: PMCPMC8463587.

26. Vigon L, Garcia-Perez J, Rodriguez-Mora S, Torres M, Mateos E, Castillo de la Osa M, et al. Impaired Antibody-Dependent Cellular Cytotoxicity in a Spanish Cohort of Patients With COVID-19 Admitted to the ICU. Front Immunol. 2021;12:742631. Epub 2021/10/08. doi: 10.3389/fimmu.2021.742631. PubMed PMID: 34616404; PubMed Central PMCID: PMCPMC8488389.

27. Tauzin A, Nayrac M, Benlarbi M, Gong SY, Gasser R, Beaudoin-Bussieres G, et al. A single dose of the SARS-CoV-2 vaccine BNT162b2 elicits Fc-mediated antibody effector functions and T cell responses. Cell Host Microbe. 2021;29(7):1137-50 e6. Epub 2021/06/17. doi: 10.1016/j.chom.2021.06.001. PubMed PMID: 34133950; PubMed Central PMCID: PMCPMC8175625.

28. Takayama K. In Vitro and Animal Models for SARS-CoV-2 research. Trends Pharmacol Sci. 2020;41(8):513-7. Epub 2020/06/20. doi: 10.1016/j.tips.2020.05.005. PubMed PMID: 32553545; PubMed Central PMCID: PMCPMC7260555.

29. Munster VJ, Feldmann F, Williamson BN, van Doremalen N, Perez-Perez L, Schulz J, et al. Respiratory disease in rhesus macaques inoculated with SARS-CoV-2. Nature. 2020;585(7824):268-72. Epub 2020/05/13. doi: 10.1038/s41586-020-2324-7. PubMed PMID: 32396922; PubMed Central PMCID: PMCPMC7486227.

30. Rockx B, Kuiken T, Herfst S, Bestebroer T, Lamers MM, Oude Munnink BB, et al. Comparative pathogenesis of COVID-19, MERS, and SARS in a nonhuman primate model. Science. 2020;368(6494):1012-5. Epub 2020/04/19. doi: 10.1126/science.abb7314. PubMed PMID: 32303590; PubMed Central PMCID: PMCPMC7164679.

31. Borakati A, Perera A, Johnson J, Sood T. Diagnostic accuracy of X-ray versus CT in COVID-19: a propensity-matched database study. BMJ Open. 2020;10(11):e042946. Epub 2020/11/08. doi: 10.1136/bmjopen-2020-042946. PubMed PMID: 33158840; PubMed Central PMCID: PMCPMC7650091.

32. Polak SB, Van Gool IC, Cohen D, von der Thusen JH, van Paassen J. A systematic review of pathological findings in COVID-19: a pathophysiological timeline and possible mechanisms of disease progression. Mod Pathol. 2020;33(11):2128-38. Epub 2020/06/24. doi: 10.1038/s41379-020-0603-3. PubMed PMID: 32572155; PubMed Central PMCID: PMCPMC7306927.

33. Bullard J, Dust K, Funk D, Strong JE, Alexander D, Garnett L, et al. Predicting Infectious Severe Acute Respiratory Syndrome Coronavirus 2 From Diagnostic Samples. Clin Infect Dis. 2020;71(10):2663-6. Epub 2020/05/23. doi: 10.1093/cid/ciaa638. PubMed PMID: 32442256; PubMed Central PMCID: PMCPMC7314198.

34. da Silva Antunes R, Pallikkuth S, Williams E, Dawen Yu E, Mateus J, Quiambao L, et al. Differential T-Cell Reactivity to Endemic Coronaviruses and SARS-CoV-2 in Community and Health Care Workers. J Infect Dis. 2021;224(1):70-80. Epub 2021/04/07. doi: 10.1093/infdis/jiab176. PubMed PMID: 33822097; PubMed Central PMCID: PMCPMC8083569.

35. Grifoni A, Weiskopf D, Ramirez SI, Mateus J, Dan JM, Moderbacher CR, et al. Targets of T Cell Responses to SARS-CoV-2 Coronavirus in Humans with COVID-19 Disease and Unexposed Individuals. Cell. 2020;181(7):1489-501 e15. Epub 2020/05/31. doi: 10.1016/j.cell.2020.05.015. PubMed PMID: 32473127; PubMed Central PMCID: PMCPMC7237901.

36. Zohar T, Alter G. Dissecting antibody-mediated protection against SARS-CoV-2. Nat Rev Immunol. 2020;20(7):392-4. Epub 2020/06/10. doi: 10.1038/s41577-020-0359-5. PubMed PMID: 32514035; PubMed Central PMCID: PMCPMC7278217.

37. Schafer A, Muecksch F, Lorenzi JCC, Leist SR, Cipolla M, Bournazos S, et al. Antibody potency, effector function, and combinations in protection and therapy for SARS-CoV-2 infection in vivo. J Exp Med. 2021;218(3). Epub 2020/11/20. doi: 10.1084/jem.20201993. PubMed PMID: 33211088; PubMed Central PMCID: PMCPMC7673958 treatment of COVID-19. D.F. Robbiani reported a patent to coronavirus antibodies pending. M.C. Nussenzweig reported a patent to anti-SARS-2 antibodies pending, and reported that Rockefeller University has applied for a patent on anti-SARS-2 antibodies. These antibodies are being produced for human clinical trials but have not been licensed to any commercial entity. No other disclosures were reported.

38. Su B, Dispinseri S, Iannone V, Zhang T, Wu H, Carapito R, et al. Update on Fc-Mediated Antibody Functions Against HIV-1 Beyond Neutralization. Front Immunol. 2019;10:2968. Epub 2020/01/11. doi: 10.3389/fimmu.2019.02968. PubMed PMID: 31921207; PubMed Central PMCID: PMCPMC6930241.

39. Beaudoin-Bussieres G, Chen Y, Ullah I, Prevost J, Tolbert WD, Symmes K, et al. A Fc-enhanced NTD-binding non-neutralizing antibody delays virus spread and synergizes with a nAb to protect mice from lethal SARS-CoV-2 infection. Cell Rep. 2022;38(7):110368. Epub 2022/02/07. doi: 10.1016/j.celrep.2022.110368. PubMed PMID: 35123652; PubMed Central PMCID: PMCPMC8786652.

40. Tauzin A, Gong SY, Beaudoin-Bussieres G, Vezina D, Gasser R, Nault L, et al. Strong humoral immune responses against SARS-CoV-2 Spike after BNT162b2 mRNA vaccination with a 16-week interval between doses. Cell Host Microbe. 2022;30(1):97-109 e5. Epub 2021/12/27. doi: 10.1016/j.chom.2021.12.004. PubMed PMID: 34953513; PubMed Central PMCID: PMCPMC8639412.

41. Ullah I, Prevost J, Ladinsky MS, Stone H, Lu M, Anand SP, et al. Live imaging of SARS-CoV-2 infection in mice reveals that neutralizing antibodies require Fc function for optimal efficacy. Immunity. 2021;54(9):2143-58 e15. Epub 2021/08/29. doi: 10.1016/j.immuni.2021.08.015. PubMed PMID: 34453881; PubMed Central PMCID: PMCPMC8372518.

42. Brunet-Ratnasingham E, Anand SP, Gantner P, Dyachenko A, Moquin-Beaudry G, Brassard N, et al. Integrated immunovirological profiling validates plasma SARS-CoV-2 RNA as an early predictor of COVID-19 mortality. Sci Adv. 2021;7(48):eabj5629. Epub 2021/11/27. doi: 10.1126/sciadv.abj5629. PubMed PMID: 34826237; PubMed Central PMCID: PMCPMC8626074.

43. Begin P, Callum J, Jamula E, Cook R, Heddle NM, Tinmouth A, et al. Convalescent plasma for hospitalized patients with COVID-19: an open-label, randomized controlled trial. Nat Med. 2021;27(11):2012-24. Epub 2021/09/11. doi: 10.1038/s41591-021-01488-2. PubMed PMID: 34504336; PubMed Central PMCID: PMCPMC8604729.

44. Wu Q, Dudley MZ, Chen X, Bai X, Dong K, Zhuang T, et al. Evaluation of the safety profile of COVID-19 vaccines: a rapid review. BMC Med. 2021;19(1):173. Epub 2021/07/29. doi: 10.1186/s12916-021-02059-5. PubMed PMID: 34315454; PubMed Central PMCID: PMCPMC8315897.

45. Munro APS, Janani L, Cornelius V, Aley PK, Babbage G, Baxter D, et al. Safety and immunogenicity of seven COVID-19 vaccines as a third dose (booster) following two doses of ChAdOx1 nCov-19 or BNT162b2 in the UK (COV-BOOST): a blinded, multicentre, randomised, controlled, phase 2 trial. Lancet. 2021;398(10318):2258-76. Epub 2021/12/06. doi: 10.1016/S0140-6736(21)02717-3. PubMed PMID: 34863358; PubMed Central PMCID: PMCPMC8639161 an investigator on studies funded or sponsored by vaccine manufacturers including AstraZeneca, GlaxoSmithKline, Janssen, Medimmune, Merck, Pfizer, Sanofi, and Valneva. She receives no personal financial payment for this work. SNF acts on behalf of University Hospital Southampton National Health Service (NHS) Foundation Trust as an investigator or providing consultative advice, or both, on clinical trials and studies of COVID-19 and other vaccines funded or sponsored by vaccine manufacturers including Janssen, Pfizer, AstraZeneca, GlaxoSmithKline, Novavax, Seqirus, Sanofi, Medimmune, Merck, and Valneva. He receives no personal financial payment for this work. ALG is named as an inventor on a patent covering use of a particular promoter construct that is often used in ChAdOx1-vectored vaccines and is incorporated in the ChAdOx1 nCoV-19 vaccine. ALG might benefit from royalty income paid to the University of Oxford from sales of this vaccine by AstraZeneca and its sublicensees under the University’s revenue sharing policy. JH has received payments for presentations for AstraZeneca, Boehringer Ingelheim, Chiesi, Ciple, and Teva. VL acts on behalf of University College London Hospitals NHS Foundation Trust as an investigator on clinical trials of COVID-19 vaccines funded or sponsored by vaccine manufacturers including Pfizer, AstraZeneca, and Valneva. He receives no personal financial payment for this work. PM acts on behalf of University Hospital Southampton NHS Foundation Trust and The Adam Practice as an investigator on studies funded or sponsored by vaccine manufacturers including AstraZeneca, GlaxoSmithKline, Novavax, Medicago, and Sanofi. He received no personal financial payment for this work. JSN-V-T is seconded to the Department of Health and Social Care, England. MR has provided post marketing surveillance reports on vaccines for Pfizer and GlaxoSmithKline for which a cost recover charge is made. MDS acts on behalf of the University of Oxford as an investigator on studies funded or sponsored by vaccine manufacturers including AstraZeneca, GlaxoSmithKline, Pfizer, Novavax, Janssen, Medimmune, and MCM vaccines. He received no personal financial payment for this work. All other authors declare no competing interests.

46. Jeyanathan M, Afkhami S, Smaill F, Miller MS, Lichty BD, Xing Z. Immunological considerations for COVID-19 vaccine strategies. Nat Rev Immunol. 2020;20(10):615-32. Epub 2020/09/06. doi: 10.1038/s41577-020-00434-6. PubMed PMID: 32887954; PubMed Central PMCID: PMCPMC7472682.

47. Wu Y, Huang X, Yuan L, Wang S, Zhang Y, Xiong H, et al. A recombinant spike protein subunit vaccine confers protective immunity against SARS-CoV-2 infection and transmission in hamsters. Sci Transl Med. 2021;13(606). Epub 2021/07/22. doi: 10.1126/scitranslmed.abg1143. PubMed PMID: 34285130.

48. Arunachalam PS, Walls AC, Golden N, Atyeo C, Fischinger S, Li C, et al. Adjuvanting a subunit COVID-19 vaccine to induce protective immunity. Nature. 2021;594(7862):253-8. Epub 2021/04/20. doi: 10.1038/s41586-021-03530-2. PubMed PMID: 33873199.

49. Wang S, Wang J, Yu X, Jiang W, Chen S, Wang R, et al. Antibody-dependent enhancement (ADE) of SARS-CoV-2 pseudoviral infection requires FcgammaRIIB and virus-antibody complex with bivalent interaction. Commun Biol. 2022;5(1):262. Epub 2022/03/26. doi: 10.1038/s42003-022-03207-0. PubMed PMID: 35332252; PubMed Central PMCID: PMCPMC8948278.

50. Junqueira C, Crespo A, Ranjbar S, de Lacerda LB, Lewandrowski M, Ingber J, et al. FcgammaR-mediated SARS-CoV-2 infection of monocytes activates inflammation. Nature. 2022. Epub 2022/04/07. doi: 10.1038/s41586-022-04702-4. PubMed PMID: 35385861.

51. Corman VM, Landt O, Kaiser M, Molenkamp R, Meijer A, Chu DK, et al. Detection of 2019 novel coronavirus (2019-nCoV) by real-time RT-PCR. Euro Surveill. 2020;25(3). Epub 2020/01/30. doi: 10.2807/1560-7917.ES.2020.25.3.2000045. PubMed PMID: 31992387; PubMed Central PMCID: PMCPMC6988269.

52. Ramakrishnan MA. Determination of 50% endpoint titer using a simple formula. World J Virol. 2016;5(2):85-6. Epub 2016/05/14. doi: 10.5501/wjv.v5.i2.85. PubMed PMID: 27175354; PubMed Central PMCID: PMCPMC4861875.

53. Mendoza EJ, Manguiat K, Wood H, Drebot M. Two Detailed Plaque Assay Protocols for the Quantification of Infectious SARS-CoV-2. Curr Protoc Microbiol. 2020;57(1):ecpmc105. Epub 2020/06/01. doi: 10.1002/cpmc.105. PubMed PMID: 32475066; PubMed Central PMCID: PMCPMC7300432.

54. Beaudoin-Bussieres G, Richard J, Prevost J, Goyette G, Finzi A. A new flow cytometry assay to measure antibody-dependent cellular cytotoxicity against SARS-CoV-2 Spike-expressing cells. STAR Protoc. 2021;2(4):100851. Epub 2021/09/21. doi: 10.1016/j.xpro.2021.100851. PubMed PMID: 34541555; PubMed Central PMCID: PMCPMC8435374.

55. Audet J, Meilleur C. Immunophenotyping for NHPs, containmnet protocol. protocolsio. 2022.

